# Collective parameter estimation of related models with an initial stability constraint

**DOI:** 10.1101/2025.11.07.684636

**Authors:** Peter Graham Clark, Lea Emmy Timpen, Alexander Heberle, Martina Prugger, Karen van Eunen, Ulrike Rehbein, Kathrin Thedieck, Daryl P. Shanley

**Affiliations:** Biosciences Institute, Newcastle University, Newcastle, UK; Institute of Biochemistry and Center for Molecular Biosciences Innsbruck, University of Innsbruck, Innsbruck, Austria; Department of Pediatrics, Section Systems Medicine of Metabolism and Signaling, University of Groningen, University Medical Center Groningen, 9713 GZ Groningen, The Netherlands; Department Metabolism, Senescence and Autophagy, Research Center One Health Ruhr, University Alliance Ruhr & University Hospital Essen, University Duisburg-Essen, Essen, Germany; German Cancer Consortium (DKTK), partner site Essen, a partnership between German Cancer Research Center (DKFZ) and University Hospital Essen, 45147 Essen, Germany; Center of Medical Biotechnology, Faculty of Biology, University of Duisburg-Essen, 45141 Essen, Germany; Westdeutsches Tumorzentrum (WTZ), 45147 Essen, Germany

## Abstract

Parameterisation of dynamic biochemical network models is a challenging aspect of systems biology. Especially when the parameter space is large and data is semi quantitive but comparable across different experimental conditions.

Here, we present a set of command line tools utilising Pycotools (COPASI) that leverages the power of high-performance computing to facilitate parameter estimation of large models with many unknown parameters. In particular, we expand upon the abilities of Pycotools to address two particular issues. Firstly, the difficulty of constraining a model’s parameterisation to assume the system begins in a steady state (prior to a perturbative stimulation). And secondly, parameterising against relative quantitative time series data that have no absolute scale.

Our software operates on the SLURM workload manager system and can be applied to any parameter estimation against time series data produced by applying a single perturbation at time zero to an equilibrated system.

We validate that our technique can produce a parameterised model of the MTOR (mechanistic target of rapamycin) network based on semi-quantitative time-series data from 2 breast cancer cell lines, stimulated with insulin and amino acids. We also show our model can make reasonable predictions on distinct signaling dynamics in one breast cancer cell line based on the other by adjusting the initial protein quantities only.

In conclusion, models should fit both the initial steady state and the dynamics following stimulation, given that stabilising systems prior to stimulation is a common experimental protocol in signaling research. By expanding standard tools, commonly used in the field, we have developed a widely applicable method, which can easily be evaluated and is amenable to wide general use.

## Introduction

Computational models based on a set of ordinary differential equations (ODEs) are frequently used to address complex mechanisms that underpin biological processes [1, 2]. Once a suitable network of interactions between components is established, a key step is to quantify parameters that govern kinetics [1, 3]. In many studies of intracellular processes, the model is fit to time series data generated from cell culture experiments. Typically, the cultures are given time to stabilise in initial reference conditions without the stimuli to be studied, the first samples are taken immediately prior to adding some stimulus, usually an agent added to the culture medium, and subsequently further samples are then taken at selected time points and measured to construct a time series dataset [4]. A model calibrated to the dataset generated in this way should ideally be constrained to begin in a stable state at the start of the simulation or at least that part of the simulation that corresponds to the period after stimulation [4, 5]. Yet, this issue is often neglected and common methods of addressing them are not always established in standard tools [4, 6].

The parameterisation of models to data under the assumption that the models start in a stable state can present a difficult challenge to modellers [4]. Many models are simply too mathematically complicated to solve for stable states analytically. When adding a lead in time the parameter estimation itself sometimes exploits selecting unstable initial conditions to achieve a close fit and longer lead in times can lead to more extreme initial conditions [4]. A variety of sophisticated methods have been developed to numerically solve both the parameter estimation optimisation problem and steady state constraint simultaneously, but these methods are often not supported or are undocumented even in popular used software, such as COPASI [4, 5]. One approach that is compatible is to incorporate a measure of the stability of initial model state into the parameter estimation objective function [7]. However, in many cases the usefulness of a non-stable state to the algorithm in generating a match to dynamic data tends to overpower the cost associated with a non-stable state in the objective function and a stable state will not be enforced.

Implementing a stable state constraint using lead in times and by applying cost functions can lead to a fitting behaviour which resists attempts to force this constraint in order to attain a better fit to dynamic data by exploiting initial conditions far from the unperturbed stable state. That was our experience at least in our first attempts to treat our data by applying a cost function to enforce an initial stable state (see S1 supplemental information, section 3, figure S2 & section 8).

Furthermore, experimental data that have been acquired with a semi quantitative technique like immunoblotting presents an additional challenge. For example, making meaningful comparisons between the levels of different proteins and phospho-proteins at different time points, or even between different experiments is only possible if samples are run on the same gel and analysed together. However, without introducing standards for each analyte (which is logistically challenging and only rarely available) it is not generally possible to infer absolute concentrations of protein or even the relative concentrations of a phospho-protein to the equivalent total protein as they are detected by distinct antibodies with different affinities to the analyte.

We have developed a method to address models resistant to conventional (initial unperturbed) stabilisation constraints in the parameter estimation process and the lack of absolute quantitative data. Approaches to handling the arbitrary units of western blot data can generally be divided into two categories: the use of scale factors as parameters and data normalisation methods [8]. Our method adapts existing software to support a previously suggested approach for data normalisation [9]. We apply the method to a test case for cell signalling networks, where we aim to explain divergent behaviour in related cell types in terms of different protein abundances in the network.

This approach is based on prior mathematical modelling indicates that different levels of protein abundance between cell lines can account for differences in the control behaviour of the same signalling pathways [10–12]. The flow of information through a network of signalling pathways can be controlled by protein abundances [13]. Changes in pathway sensitivity over time and between cells at the same time may also be explained by variations in protein abundance [14]. Mathematical modelling has also addressed cytoplasmic domain binding competition on cell surface receptors [15]. This indicates that protein abundance can effectively act as a junction box ‘wiring’ cell surface receptors into different pathways.

This is the problem that arises in the context of related immortalised cell lines [10]. Also cancer cells of the same type from different patients can have quite different behaviour [11]. In some cases it may be possible to explain this divergent behaviour in terms of variations in protein abundance [10, 11]. Here, we investigate two different breast cancer cell lines treated with insulin and amino acids to monitor the MTOR signalling pathway. We use this to demonstrate the method can parameterise models that can describe the behaviour of multiple cell lines and also infer the behaviour of cell lines not used in its parameterisation.

## Materials and methods

### Approach

Our basic approach can be summarised as adjusting a user defined biochemical network model, which we term a base model, to programmatically include a pre-processing and post-processing step. The pre-processing step uses analytical methods to estimate initial conditions (IC) close to a stable state then runs the model until a stability test is satisfied. Only then is the appropriate stimulation applied and the desired time course simulated. Experimental time points are measured relative to this stimulation. This represents what we call the intermediate model. Multiple copies of this intermediate model (representing different cell lines or preconditioning protocols) are incorporated into the final model that performs post processing. Incorporation of multiple sub models is achieved using a sub model feature in the associated software similar to the way a function can be imported and reused in code. Once each copy of the intermediate model has finished acquiring the data for all time points the post processing step calculates versions of each species normalised over time points and model copies (which in our case represented different cell lines, i.e. MCF-7 and ZR-75-1, but could also relate e.g. to different preconditioning protocols) which are used for comparison with experimental data in the parameter estimation (Fig 1).

**Figure 1.**
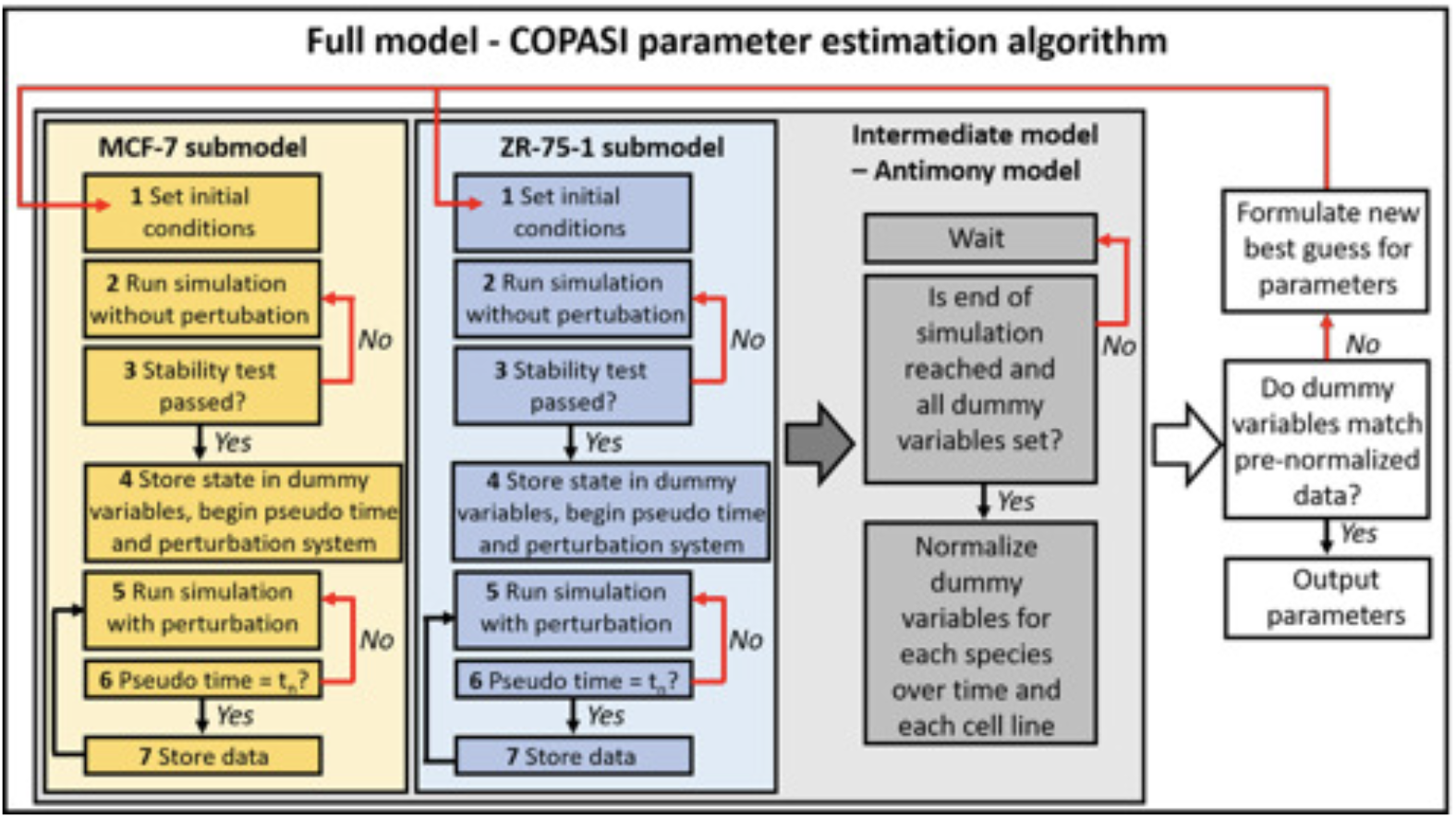
Example scheme of the workflow. The two submodels for data from MCF-7 and ZR-75-1 cells (created by stringWangler.py) are incorporated into an intermediate model (an Antimony string created by modelParaliser.py), which is run within the COPASI parameter estimation algorithm (by GPParamiteriser.py). The workflow is conducted in the following steps; 1. The initial conditions are set based on the parameters and protein abundance. 2. Simulation is run without perturbation. 3. Stability test is performed. If test fails, step 2 is repeated. 4. When stability test is passed dummy variables for *t*_0_ are assigned, perturbation is applied to the system and the pseudo time timer is started. 5. Simulations are run with perturbations. 6. Test if pseudo time is reached *t*_*n*_. If not step 5 is repeated. 7. If yes, the data for this time point is stored. Step 5 to 7 is repeated until all time points are simulated and stored. The intermediate model waits until simulations are completed and all the dummy variables are set. If this does not occur a very large number is returned in all dummy variables. If it does occur the dummy variables are normalized for each species over time and for each cell line. Then, the COPASI parameter estimation algorithm compares the dummy variables to the pre-normalized experimental data. If the data match, the algorithm gives the output parameters. When this is not the case, the algorithm formulates a new best guess for the starting parameters and the procedure begins again in step 1.

In this way all the information about the absolute scale of any species is removed from the outputs of the model but information about variation over time and between cell lines (or preconditioning steps) is retained. The model is constructed in such a way that if stabilisation takes too long and there is insufficient time to complete pre or post processing steps the outputs delivered to the external back-end parameter estimation algorithm are kept at high levels to ensure the rejection of the parameter set associated with the simulation. In this way, any parameter sets that fail to quickly achieve stability are likely to be discarded by the algorithm.

A common format for system biology models is Systems Biology Markup Language (SBML) [16, 17], however, for constructing the intermediate and final model it is more convenient to use Antimony, an alternate format with portability to and from SBML [18]. We have created 2 command line tools, one (stringWangler.py) to create the intermediate and one (modelParaliser.py) to create the final model. The first (stringWangler.py) derives and inserts initial conditions based on parameters and provides for a variable length stabilization lead in period. To do this it parses the Antimony string defining the base model converting its reactions to a set of Sympy expressions for the equivalent ODEs. Sympy is a widely used python based computational algebra library [19]. This generates equations for the stable state. Options are provided to perform a number of user directed substitutions, solutions and simplifications that will give a set of an analytical guesses for most but not necessarily all the initial conditions in the model. Initial conditions set in this way will begin with a zero time derivative for any choice of parameters.

For example, to set the initial conditions for the system of equations found in Table S4 all derivatives are set to 0 and the pre-perturbation conditions substituted in. Then as indicated by the configuration file (stabAdj2.json), stringWangler.py will first substitute out the variables RPS6KB1, IRS1, AKT1, TSC2-pT1462, EIF4EBP1 and AKT1S1-pT246 using the appropriate conservation equations. The associated stability equations are ignored. Then as directed by the configuration file it will attempt to solve IRS1-pS636/639’s stability equation throwing an error if it does not find a unique solution. This expression for IRS1-pS636/639 will be substituted into the remaining equations. The process proceeds in the same manner with the remaining equations in the order instructed by the configuration file. We use sympy to perform these steps within our software.

This reduces the amount of ‘lead in’ time our model needs to reach its stable state (see supplementary S1.5). The code generates the appropriate lines for setting these initial conditions and inserts them into the Antimony string (facilitating step 1 Fig 1).

A further algorithm calculates physically plausible initial condition estimates for remaining species based on the total availability levels of the various proteins involved. Lines setting these initial conditions are also inserted (facilitating step 1 Fig 1).

The code also creates a new reaction with a new dummy species (facilitating step 4 & 6 Fig 1). This species is effectively a pseudo time measurement. The reaction is constructed so that this species will begin at zero and only when a stability test as defined by an expression based on the ODEs are satisfied will it start increasing a constant rate of 1 per unit of time (see Eq 1). To achieve this an event statement is constructed and inserted into the Antimony string (facilitating step 3 Fig 1) which takes as its trigger conditions that the absolute value of each time derivative (of each species *x*_*i*_) in the original model should be below a very small number (*ε*_*i*_) (see Eq 2). It’s also necessary to have as a condition that the pseudo time parameter be zero to ensure this event is only triggered once.

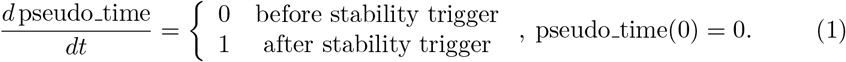

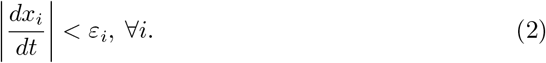

The event has 3 effects (in step 4 Fig 1). It triggers the stimulation that begins the section of simulation, which corresponds to the experimental measurements (step 5 of Fig 1). It changes the parameter associated with the pseudo time from 0 to 1 causing pseudo time to increase acting as a measure of how much time has passed in the simulation since stimulation. Lastly it stores the value of the time = 0 timepoint in a set of dummy variables. Additional events are added to store other timepoints in dummy variables using the pseudo time species as a trigger (facilitating step 6 & 7 in Fig 1). As previously mentioned, these dummy variables are initialised to very large numbers.

GPParamiteriser.py uses the parameter estimation algorithm built into the systems biology COPASI application [20]. In COPASI our models have only one time point at the end of the simulation from which the dummy variables can be read. Making comparisons at one time point is problematic as by default the importance of species in data are assigned weights in COPASI. To overcome this issue our code uses the Pycotools3 package to interface with COPASI [21]. To disable this weighting behavior it’s necessary to set the “Normalize Weights per Experiment” option to “standard_deviation” which in Pycotools is the same as setting “weight_method” to “standard_deviation”. This will ensure that the weights on the data are all one.

The second stage uses the intermediate model (as defined by the antimony file created by the 1st phase) to create the one upon which the parameter estimation will be run. We again created a python command line tool (modelParaliser.py) to automate this process as much as possible. The code takes advantage of the modularity provided by the Antimony language. The intermediate model is imported as a sub model (a formal definition of a model structure in Antimony, multiple copies of which can be incorporated into larger models) and the main model defines as many instances of this sub model as needed for each cell line. Parameters that are to be estimated or set identically between the sub models are linked to new parameters with the same name in the new model. In the event that multiple experimental conditions are to be addressed the new model can override initial conditions in the sub models to represent differing culture conditions or perturbations. Parameters in the sub model that represent total protein amounts (see Application section) from which initial conditions are calculated or that operate as dummy variables are linked to separate parameters in the new model with suffixes indicating the relevant sub model. Additionally, the pseudo time species for the sub models are linked to distinct variables in the new model. The model needs to reference them to ensure that post processing calculations only take place after each sub model simulation has completed gathering data.

This new model has 2 new events. The first is triggered one unit of time after all sub models have acquired their last time point and calculates the average value over time and over each sub model of each species as given by the relevant dummy parameters. The second event takes place one unit of time later and uses these averages, provided they are non zero, to divide through the dummy values normalizing them. This is necessary because COPASI doesn’t have a documented option to automatically normalise simulation data within parameter estimations. The experimental data passed to COPASI must be scaled to account for this normalisation. As with the intermediate model all dummy parameters have their initial values set to very high numbers as a guard against late or non-stabilisation.

The resulting antimony string is used by GPParamiteriser.py which, when run on a SLURM computer cluster, passes it to many instances of COPASI running in parallel. Within COPASI the submodels run their simulations, the intermediate model checks if the simulations are completed and if all the dummy variables are set. Thereafter, it will normalize the dummy variables for each species (in our case for each protein or phosphoprotein) over time for each cell line (in our case MCF-7 and ZR-75-1). Then the COPASI parameter estimation algorithm will check if the dummy variables fit to the pre-normalized data and gives as an output the parameters. When the data does not match, the COPASI parameter estimation algorithm will create a new best guess for the parameters. The results are returned to GPParamiteriser.py and stored for analysis.

### Application

We demonstrate this method with immunoblot data used to parameterise a model to two breast cancer cell lines, MCF-7 and ZR-75-1 (Fig 2). As will be shown, our intention here was to test whether it is possible to use our method to construct and parameterise a model that can describe the behaviour of the MTORC1 signalling network in several breast cancer cell lines only by modifying its initial conditions [1]. The experience of our research group is that parameter estimations will often push initial conditions into nonstable states, despite constraints to achieve initial stability. Presumably this is because the process prioritises a good match to dynamic data that is difficult to fit. Enforcing an initial stable state in a difficult parameter estimation was the vital obstacle to overcome in this work. The cell line signalling pathways were assumed to start in a stable state before being subjected to a stimulus (insulin and amino acids) at the first time point. The model was not amenable to an analytical solution of the stable state and lead in periods proved ineffective.

**Figure 2.**
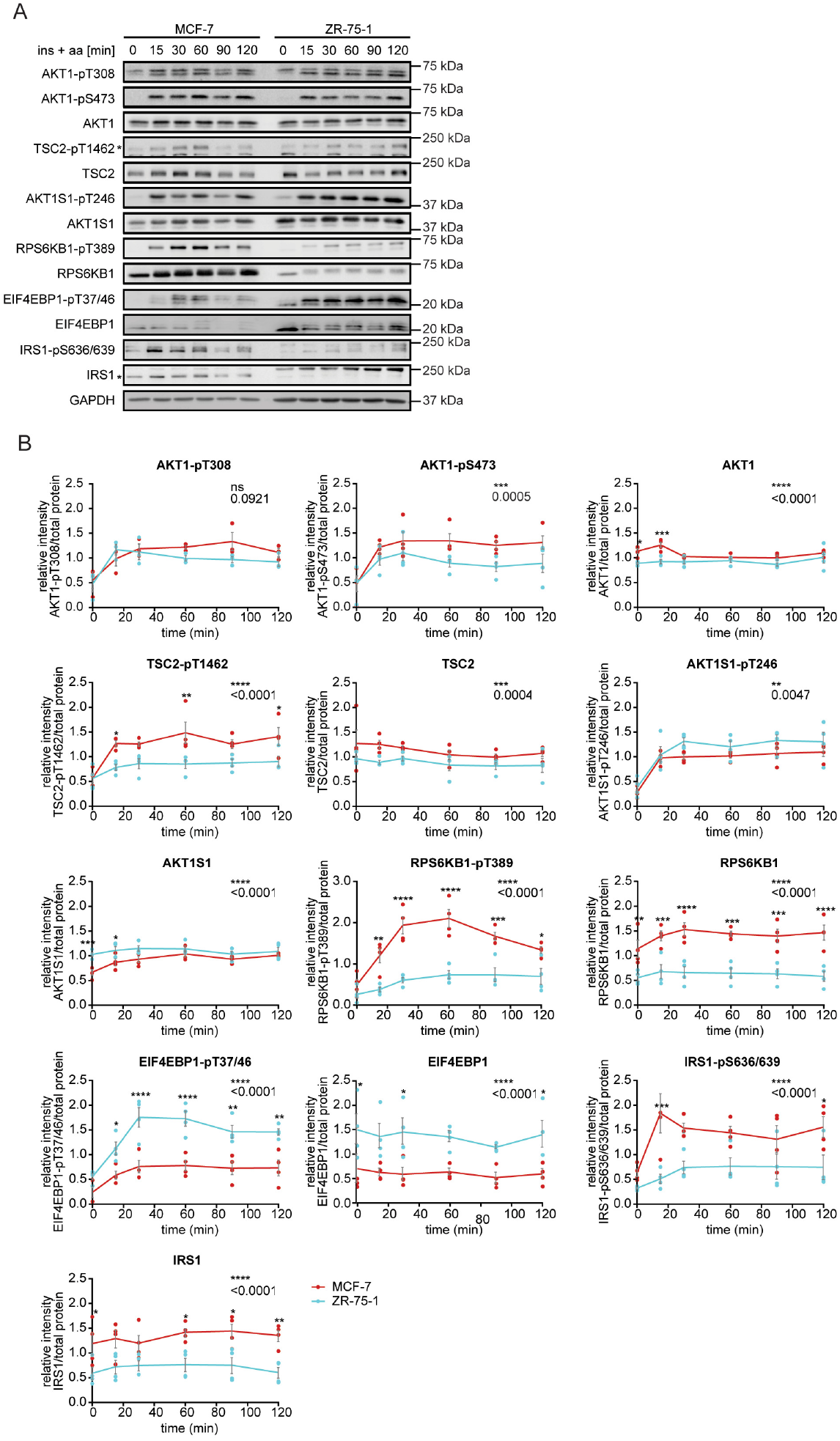
Stimulation time course data in two different breast cancer cell lines. **(A)** MTOR signalling dynamics in MCF-7 and ZR-75-1 cells. Cells were stimulated with insulin and amino acids for the indicated time points. AKT1-pT308, AKT1-pS473, TSC2-pT1462, AKT1S1-pT246, RPS6KB1-pT389, EIF4EBP1-pT37/46, IRS1-pS636/639 and their respective total protein levels were monitored by immunoblot. Data represent 4 biological replicates. For IRS1 and TSC2-pT1462 * indicates the corresponding protein band. **(B)** Quantification of data shown in (A). Line shows the mean values. Dots show individual data points. The MCF-7 and ZR-75-1 cell lines were compared using a two-way ANOVA followed by a Bonferroni multiple comparison test. n = 4 independent biological replicates. Shown are single data points ±SEM. ns=not-significant; *\** p ≤0.05; *p≤0.01; * * * p ≤0.001; * * ** p≤0.0001. Abbreviations are defined in the text. (Quantitations of each replicate, as well as the mean are shown in the supplements S2-4.)

Likewise scales to convert immunoblot data to absolute concentrations cannot be readily established and at best required an iterative process that required many repeated parameter estimations. We developed two models of the relevant signalling networks, one including MTOR dynamics (Fig 3) and a simplified version without MTOR (Fig 4), which were coded into Antimony strings and then converted into meta models containing either an MCF-7 and ZR-75-1 sub model or models containing both sub models. These meta models were used for parameter estimation and by design withhold stimulation until the model has stabilised and ensures comparisons are made relative to the moment of stabilisation.

**Figure 3.**
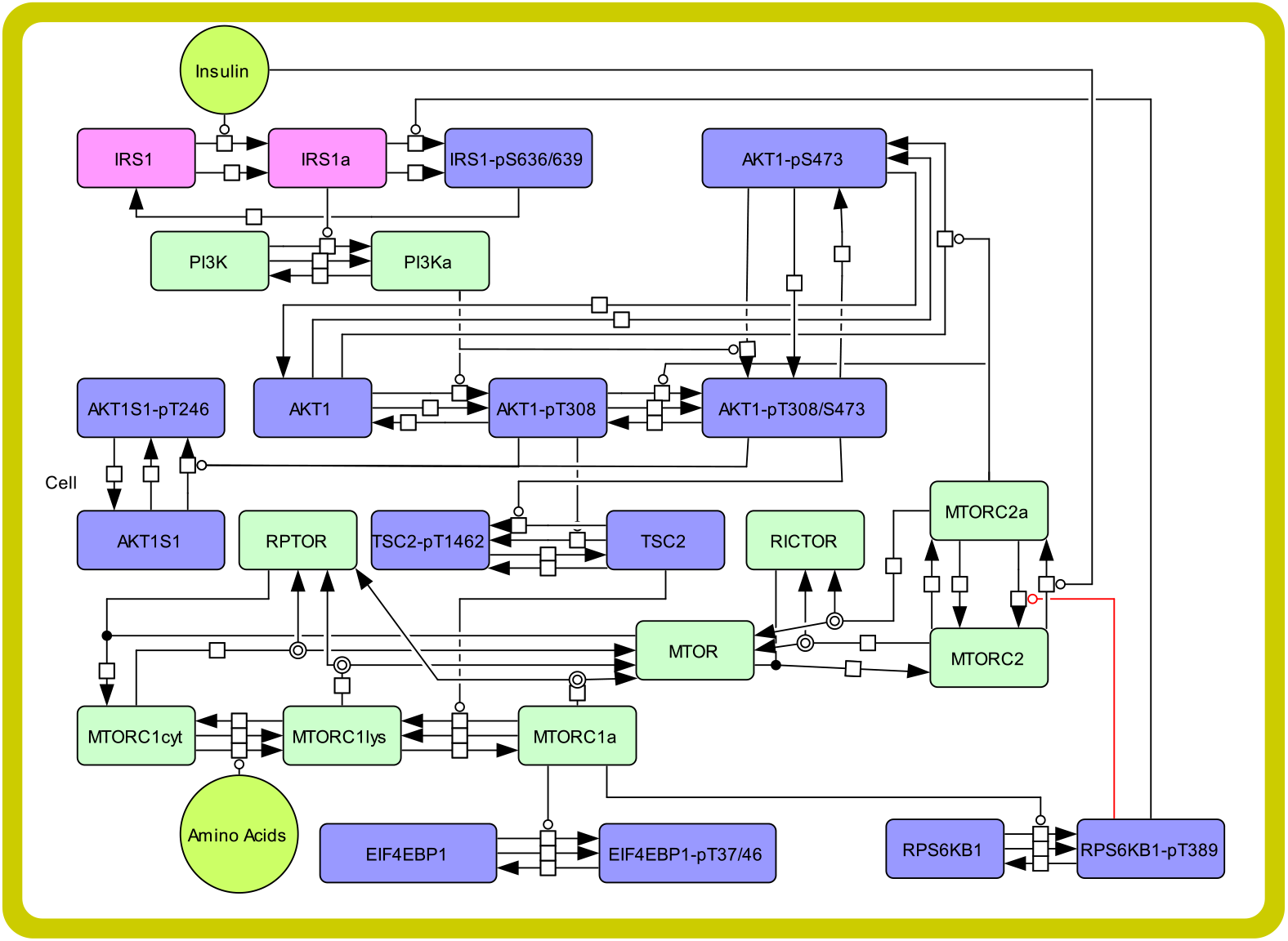
Structure of MTOR model. Purple boxes represent species for which semiquantitative immunoblot data for both the total protein and phospho-protein levels were available. For IRS1 (pink) semiquantitative immunoblot data for the total level and phosphorylation at serine 636/639 were available. The phosphorylation was indicative of inhibited IRS in response to active RPS6KB1 but not of activated IRS1 (IRS1a, pink) in response to insulin. Light green boxes represent species for which we had no direct measurements and lime green circles the stimulating factors used to perturb the model. All abbreviations and protein names are defined in the text. (Equations are shown in the supplement S1.)

**Figure 4.**
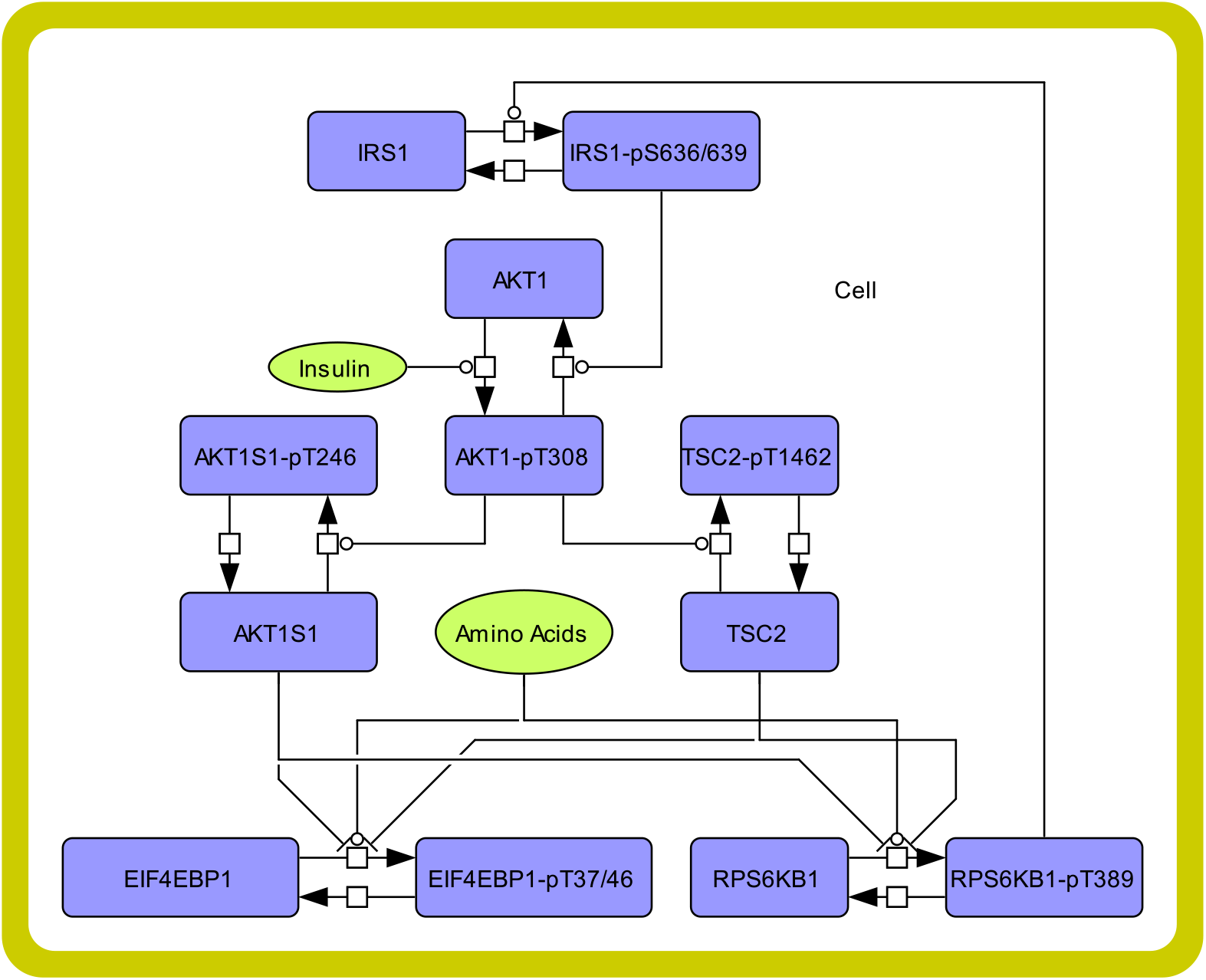
Structure of reduced model. Blue boxes represent species for which semi quantitative immunoblot data for both the total protein and phospho-protein levels were available. Lime green circles represent the stimulating factors used to perturb the model. Protein names and abbreviations are defined in the text. (Equations are shown in the supplement S1.)

Once the data for comparison has been acquired for all sub models of the meta model the post processing step calculated normalised over time and cell lines versions of each species (Fig 1).

### Cell culture and stimulation protocols

Experiments were performed using the MCF-7 and ZR-75-1 cell lines that were obtained from ATCC (HTB-22, CRL-1500). Cells were maintained at a humidified atmosphere of 37 °C and 5% CO2 in Dulbecco’s Modified Eagle Medium with 4.5 g/L glucose (DMEM; PAN Biotech, Cat. No. P0403600) supplemented with 10% fetal calf serum (FCS; Gibco, Life Technologies, Cat. No. 10270-106) and 3 mM L-glutamine (Gibco, Life Technologies, Cat. No. 25030-024). Prior to the time-course experiments, cells were washed twice with phosphate-buffered saline (PBS; PAN biotech, Cat. No. P04-36500) and 16 hours starved for amino acid and growth factors in Hank’s buffered salt solution (HBSS; PAN Biotech, Cat. No. P04-32505). The cells were washed once with PBS and stimulated for the indicated time periods with DMEM with 1 g/L glucose (pan-biotech, Cat. No. P04-01500), 3 mM L-glutamine and 100 nM insulin (Sigma-Aldrich, Cat. No. I1882).

### Antibodies and reagents

The following antibodies were purchased from Cell Signaling Technology: AKT1 (Cat. No. 4691), AKT1-pT308 (Cat. No. 2965), AKT1-pS473 (Cat. No. 4060), TSC2 (Cat. No. 3635), TSC2-pT1462 (Cat. No. 3617), AKT1S1 (Cat. No. 2691), AKT1S1-pT246 (Cat. No. 2997), RPS6KB1 (Cat. No. 2708), RPS6KB1-pT389 (Cat. No. 9206), EIF4EBP1 (Cat. No. 6459), EIF4EBP1-pT37/46 (Cat. No. 9452), IRS1 (Cat. No. 2382), IRS1-pS636/639 (Cat. No. 2388). GAPDH (Cat. No. ab8245) was purchased from Abcam. All antibodies were diluted 1:1000, except for GADPH, which was diluted 1:10,000. Primary antibodies were diluted in tris-buffered saline-Tween (TBST; 138 mM sodium chloride, 2.7 mM potassium chloride, 66 mM TRIS base [pH 7.4], 0.1% Tween-20), supplemented with 5% bovine serum albumin (BSA; Sigma-Aldrich, Cat. No. A2153) and 0.1% NaN3. Horseradish peroxidase-conjugated goat anti-mouse (Cat. No. 31430) and goat anti-rabbit IgG (Cat. No. 31460) were purchased from Thermo Scientific Pierce and were diluted 1:4000 in TBST containing 2.5% BSA.

### Cell lysis and immunoblotting

For cell lysis, cells were washed twice with PBS and lysed with radio immunoprecipitation assay (RIPA) buffer (1% IGEPAL CA-630, 0.1% sodium dodecyl sulfate (SDS), and 0.5% sodium deoxycholate in PBS), supplemented with Complete Protease Inhibitor Cocktail (Sigma-Aldrich, Cat. No. 11836145001), Phosphatase Inhibitor Cocktail 2 (Sigma-Aldrich, Cat. No. P5726) and Cocktail 3 (Sigma-Aldrich, Cat. No. P0044). Protein Assay Dye Reagent (Bio-Rad, Cat. No. 500-0006) was used to measure protein concentrations and samples were adjusted to the lowest value.

Samples were mixed with sample buffer (10% glycerol, 1% β-mercaptoethanol, 1.7% SDS, 62.5 mM TRIS base [pH 6.8], bromophenol blue) and heated for 5 min at 95 °C to denature the proteins. Proteins were separated by size using SDS polyacrylamide gel electrophoresis (SDS-PAGE). Dependent on the protein size, the acrylamide percentage ranged between 8 to 14%. For the separation, 90 to 190 V was applied to a Mini-PROTEAN Tetra Vertical Electrophoresis Cell system (Bio-Rad, Cat. No. 1658029FC) containing running buffer (0.2 M glycine, 25 mM TRIS base, 0.1% SDS). For total protein staining, gel was washed in MilliQ water and stained with SimplyBlue Safestain (Invitrogen, Cat. No. LC6060) overnight. After destaining with MilliQ water, images were taken with using the LAS-4000 mini camera system (GE Healthcare). For protein transfer, the Mini Trans-Blot Electrophoretic Transfer Cell (Bio-Rad, 1703930) was filled with transfer buffer (100 mM glycine, 50 mM TRIS base, 0.01% SDS, [pH 8.3], 10 % methanol). 45 V was applied for 1 hour and 50 minutes to transfer the proteins to polyvinylidene difluoride (PVDF) membranes (Merck, Cat. No. IPVH00010).

Membranes were blocked with 5% BSA in TBST for 20 minutes and primary antibodies were incubated overnight at 4 °C. Membranes were washed with TBST and incubated with HRP-coupled secondary antibody (goat-anti-mouse, goat-anti-rabbit) for at least two hours. Membranes were washed again with TBST and Pierce ECL western blotting substrate (Thermo Fisher Scientific, Cat. No. 32209) or SuperSignal West FEMTO (Thermo Fisher Scientific, Cat. No. 34095) was added for detecting chemiluminescence using a LAS-4000 mini camera system (GE Healthcare). Raw images were converted to red green blue (RGB) color and exported as TIFF files using ImageJ and further processed with Adobe Photoshop CS6. Quantifications of immunoblots were obtained by using ImageQuant TL (GE healthcare). Background substraction from the raw files was performed with the Rolling Ball method, with a defined radius of 200 for all images. The pixel values were normalized to the average pixel intensity of all conditions and was then normalized to the total protein amount. Statistical analysis was performed in GraphPad Prism 9. Two-way-ANOVA test was used for comparing the different cell lines over the complete time-course, followed by Bonferroni’s multiple comparison test, to analyse the significance of the single time-points.

### MTOR Model

The model selected to fit this data is shown in Fig 3. This model is derived from a series of previously established models [6, 23–25] and is suitably complex model for our purpose with all reactions defined by mass action kinetics.

### Reduced Model

A reduced version of the MTOR model was created in which the stable state could be solved analytically (Fig 4). The ODEs associated with the reduced model give rise to a set of stationary state equations that simplify to the solution of a cubic equation. We verified this ‘by hand.’ We demonstrate that our method successfully reparameterises this model as a proof of concept.

### Parameter estimation

Parameter estimation was performed with custom python code using Pycotools and COPASI on a SLURM based compute cluster (the Newcastle university rocket HPC cluster). Parameter estimation was a global chaser type composed of a particle swarm with a swarm size of 100 and an iteration limit of 4500 followed by a Hooke Jeeves parameter estimation with an iteration limit of 50, tolerance of 0.00001 and rho of 0.2. 200 copies of the parameter estimation were run (on 200 cores that could require up to 2 days of compute time on each) and the best 10, as indicated by RSS scores, were simulated to observe their fit to data. The model was given an additional 202 minutes of simulated time within the parameter estimation. 200 to facilitate stabilisation prior to the simulation proper and two for the 2-time units the model uses for post simulation calculations.

Experimental data was converted to a single time point of data for the set of dummy parameters as described previously (in the approach section). The total protein levels for AKT1S1, AKT1, EIF4EBP1, RPS6KB1, TSC2 and IRS1 were calculated from experimental data and passed to the parameter estimation as independent variables. The total protein levels for MTOR, RICTOR, RPTOR and PI3K were estimated along with the model parameters.

To facilitate the examination, the 10 best parameter sets (as indicated by RSS) were simulated on the original model for both cell lines. Because phosho-protein levels in the model have an absolute scale but the phospho-protein levels in experimental data do not, to facilitate generation of figures for comparison a scale factor was applied to the simulated data to match its average level (the mean over the appropriate time points in both cell lines for each measurable) to that of the experimental data.

A custom ‘goodness of fit’ algorithm (a simple sum of squares for each compared set of post scaled data points) that recalculates an RSS score was used to select the best cases out of the 10-time courses returned (as indicated in figures).

### Data and Code Repository

Data and code for this work can be found at https://github.com/npc101-ncl/MESI_PLOS_COMP_BIO.

### Nomenclature

For proteins, HUGO names and symbols are used in the text. In the code adapted names are used, which were based on commonly used names (Table 1).

**Table 1.** Protein names.

| HUGO protein abbreviation | HUGO full protein name | Common name | Abbreviation in code |
| --- | --- | --- | --- |
| AKT1 | AKT serine/threonine kinase 1 | Akt | Akt |
| AKT1-pS473 | AKT serine/threonine kinase 1 phosphorylated at serine 473 | Akt-pS473 | AktpS473 |
| AKT1-pT308 | AKT serine/threonine kinase 1 phosphorylated at threonine 308 | Akt-pT308 | AktpT308 |
| AKT1S1 | AKT1 substrate 1 | PRAS40 | PRAS40 |
| AKT1S1-pT246 | AKT1 substrate 1 phosphorylated at threonine 246 | PRAS40-pT246 | PRAS40pT246 |
| EIF4EBP1 | Eukaryotic translation initiation factor 4E binding protein 1 | 4E-BP1 | FourEBP1 |
| EIF4EBP1-pT37/46 | Eukaryotic translation initiation factor 4E binding protein 1 phosphorylated at threonine 37 and 46 | 4E-BP1-pT37/46 | FourEBP1pT37_46 |
| IRS1 | Insulin receptor substrate 1 | IRS1 | IRS1 |
| IRS1a | active form of IRS1 | – | IRS1a |
| IRS1-pS636/639 | Insulin receptor substrate 1 phosphorylated at serine 636 and 639 | IRS1-pS636/639 | IRS1pS636_639 |
| MTOR | Mechanistic target of rapamycin kinase | mTOR | mTOR |
| MTORC1 | MTOR complex 1 | mTORC1 | mTORC1 |
| MTORC1a | active form of MTORC1 | – | pmTORC1 |
| MTORC2 | MTOR complex 2 | mTORC2 | mTORC2 |
| MTORC2a | active form of MTORC2 | – | pmTORC2 |
| PI3K | phosphoinositide 3-kinase | PI3K | PI3K |
| RICTOR | RPTOR independent companion of MTOR complex 2 | Rictor | RICTOR |
| RPS6KB1 | ribosomal protein S6 kinase B1 | S6K | S6K |
| RPS6KB1-pT389 | ribosomal protein S6 kinase B1 phosphorylated at threonine 389 | S6K-pT389 | S6KpT389 |
| RPTOR | regulatory associated protein of MTOR complex 1 | Raptor | RPTOR |
| TSC2 | TSC complex subunit 2 | TSC2 | TSC2 |
| TSC2-pT1462 | TSC complex subunit 2 phosphorylated at threonine 1462 | TSC2-pT1462 | TSC2pT1462 |

## Results

### MTOR signaling in breast cancer cell lines

To test whether it is possible to construct and parameterise a model that can describe the behaviour of different cell lines only by modifying its initial conditions, we focused on MTOR signalling in the two breast cancer cell lines, MCF-7 and ZR-75-1.

MTOR is a conserved serine/threonine kinase which resides in two multiprotein complexes, termed MTOR complex 1 (MTORC1) and 2 (MTORC2) [1]. Being a central regulator of cellular growth and metabolism, alterations of this network are important determinants of ageing and age-related diseases, including cancer. While mTORC1 and MTORC2 share the same catalytic subunit, MTORC1 contains the specific interactors regulatory associated protein of MTORC1 (RPTOR, also named Raptor) and AKT1 substrate 1 (AKT1S1, also named PRAS40). MTORC2 specifically contains RPTOR independent companion of MTORC2 (RICTOR) and MAPK associated protein 1 (MAPKAP1) [26–35]. We focused our model on the network encompassing MTORC1 and comprising numerous feedback loops.

The MTORC1 signalling network is constructed through the interplay of kinases, which either activate or inhibit each other. A protein may undergo phosphorylation at one or multiple sites, dictating its functionality. The MTOR network topology and signal transition have been reviewed in detail by Heberle et al [1, 36]. In brief, activation of the MTORC1 network is regulated by external cues, including growth factors such as insulin. Binding of insulin to the insulin receptor leads to the recruitment and activation of insulin receptor substrate (IRS1), triggering membrane translocation and activation of phosphoinositide 3-kinases (PI3K). Downstream of PI3K, AKT serine/threonine kinase 1 (AKT1) is phosphorylated at threonine 308 (AKT1-pT308). Phosphorylation of AKT1 at serine 473, however, is mediated by MTORC2. Both phosphorylation events activate AKT1. When phosphorylated at the T308 site, AKT1 negatively regulates the MTORC1 inhibitory proteins AKT1S1 by phosphorylation at threonine 246 (AKT1S1-pT246) and TSC complex subunit 2 (TSC2) through phosphorylation at threonine 1462 (TSC2-pT1462). Under amino acid replete conditions, MTORC1 translocates from the cytosol (MTORC1cyt) to the lysosomal membrane (MTORC1lys). MTORC1 phosphorylates a variety of targets including eukaryotic translation initiation factor 4E binding protein 1 (EIF4EBP1) at threonine 37 and 46 (EIF4EBP1-pT37/46) and ribosomal protein S6 kinase B1 (RPS6KB1) at threonine 389 (RPS6KB1-pT389). RPS6KB1 in turn inhibits IRS1 by phosphorylation at serine 636 and 639 (IRS1-pS636/639), this mediating negative feedback to upstream insulin signalling.

For model parameterisation, we monitored the levels of 6 key proteins and 7 phosphorylation states across the MTOR network in the two cell lines. Dynamic time course data were recorded upon stimulation with insulin and amino acids. Growth factors and amino acids stimulation increased phosphorylation levels throughout the MTOR signalling pathway, whereas the total protein levels stayed the same (Fig 2). MCF-7 cells exhibited higher protein and phosphorylation levels than ZR-75-1 cells, except for AKT1-pT308, AKT1S1 total and AKT1S1-pT246, and the MTORC1 substrate EIF4EBP1 total and EIF4EBP1-pT36/46. AKT1-pT308 was not altered between the two cell lines. For each cell line and model combination we performed the parameterisation as previously described. Of the 10 best parameter sets (per cell line and model as indicated by Pycotools RSS score) we selected the one (for each model and cell line) with the best fitness output scores as generated by our visualisation code.

When we attempted to parameterise the full MTOR network model (Fig 3) to MCF-7 data alone the simulations were consistent with data (Fig 5).

**Figure 5.**
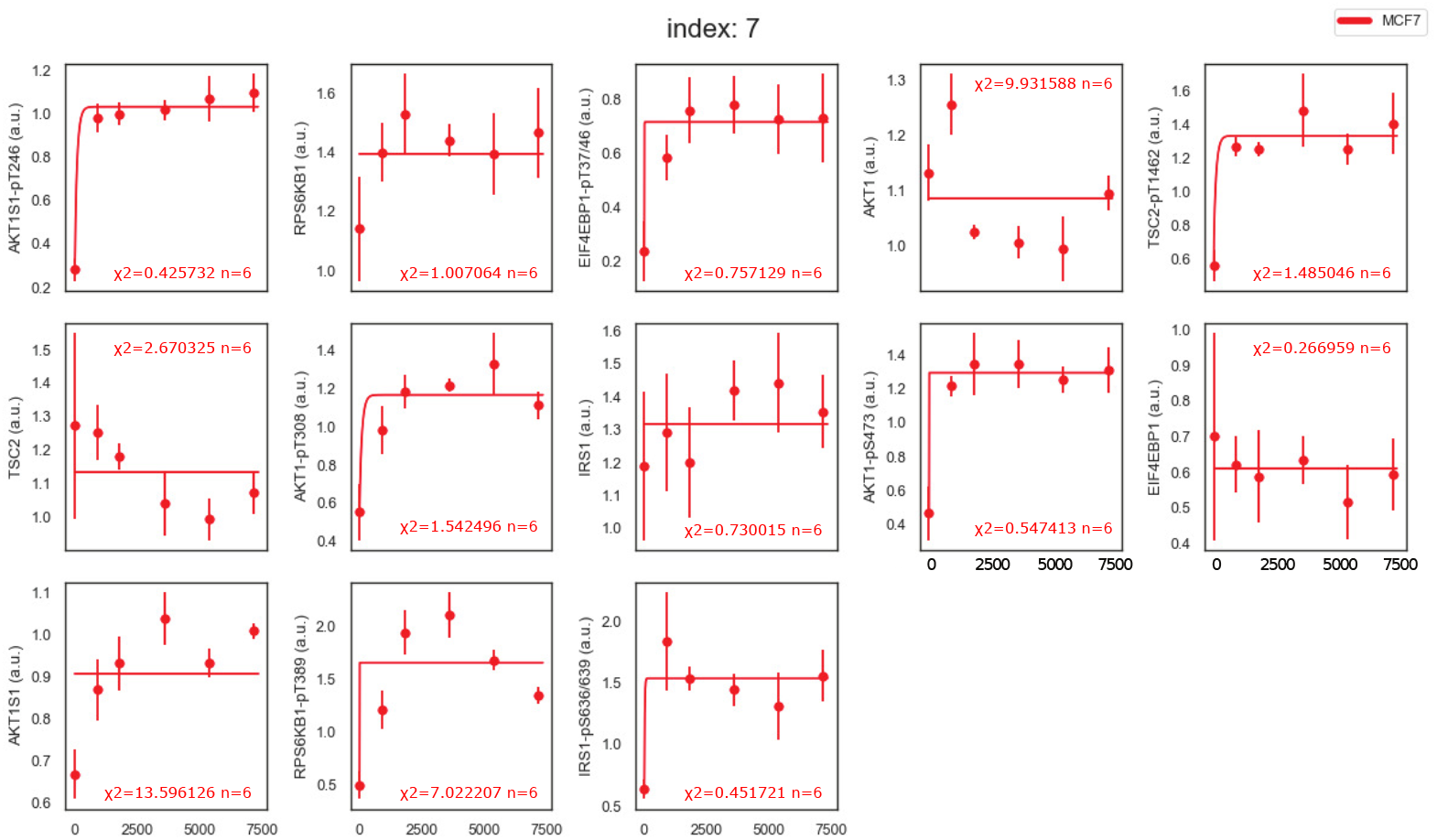
MCF-7 based parameterisation of TOR model. MTOR model parameterised against MCF-7 data (see supplement S3). Parameter set 7 out of sets 0 to 9 which closely matches data (chosen based on custom RSS score). Note that each phospho-protein species (AKT1S1-pT246, AKT1-pS473, AKT1-pT308, TSC2-pT1462, RPS6KB1-pT389, EIF4EBP1-pT37/46 and IRS1-pS636/639) has had a scale factor applied to the simulation outputs to match them to the experimental data (which has no absolute quantification). Total protein levels (AKT1, AKT1S1, TSC2, RPS6KB1, EIF4EBP1 and IRS1) were not scaled. It should be noted AKT1-pT308/S473 levels contribute to both the AKT1-pS473 and AKT-pT308 readings. *χ*^2^ values as described in supplementary S1.4. Dots, experimental data; lines, simulation; bars, standard deviation.

Likewise, when we parameterised our model incorporating the full MTOR network to both MCF-7 and ZR-75-1 data simultaneously we observed a close match to experimental data (see Fig 6). Similar to the experimental data we observed higher levels of phosphorylation levels in MCF-7 cells, except for EIF4EBP1 and AKT1S1. However, in initial attempts to reparameterise the MCF-7 model using protein levels from ZR-75-1 cells, and vice versa, the simulations were inconsistent with data.

**Figure 6.**
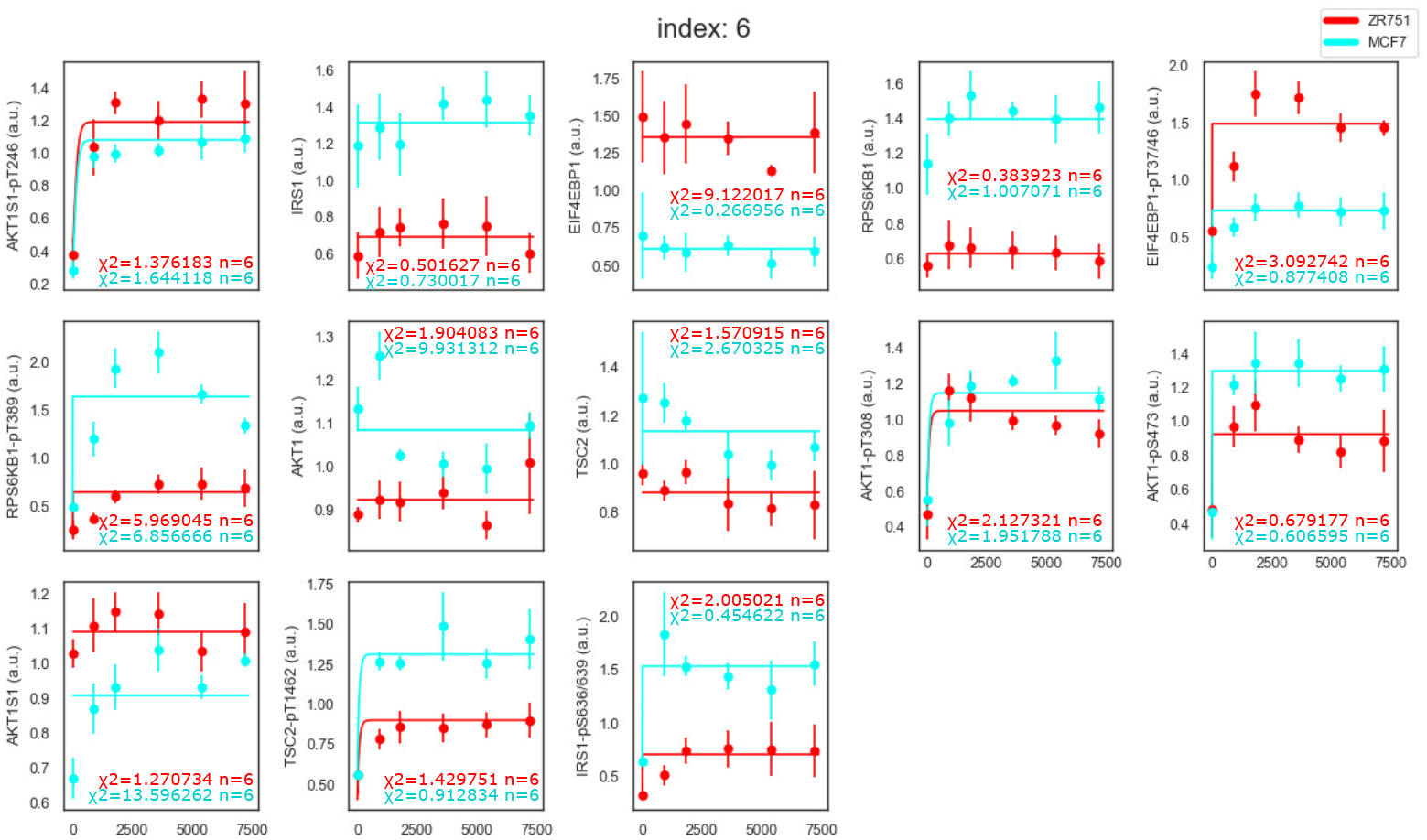
MCF-7 & ZR-75-1 based parameterisation of MTOR model. MTOR model parameterised against MCF-7 and ZR-75-1 data (see supplement S2). Parameter set 6 out of sets 0 to 9 which closely matches data (chosen based on custom RSS score). Note that each phospho-protein species (AKT1S1-pT246, AKT1-pS473, AKT1-pT308, TSC2-pT1462, RPS6KB1-pT389, EIF4EBP1-pT37/46 and IRS1-pS636/639) has had a scale factor applied (identical in both cell lines) to the simulation outputs to match them to the experimental data (which has no absolute quantification). Total protein levels (AKT1, AKT1S1, TSC2, RPS6KB1, EIF4EBP1 and IRS1) were not scaled. It should be noted AKT1-pT308/S473 levels contribute to both the AKT1-pS473 and AKT1-pT308 readings. *χ*^2^ values as described in supplementary S1.4. Dots, experimental data; lines, simulation; bars, standard deviation.

We considered that reducing the number of parameters may improve the fit between simulations and data when reparameterising a model on protein levels from a different cell line. We opted for our reduced model (Fig 4) to not include MTORC1 itself, but only its upstream cue AKT1 that phosphorylates the two MTORC1 inhibitory proteins AKT1S1 and TSC2, and the MTORC1 substrates EIF4EBP1 and RPS6KB1, as well as IRS1 as a mediator of negative feedback from RPS6KB1 to AKT1. Insulin activates AKT1 whereas amino acids impinge on MTORC1 downstream of AKT1 and TSC2, and were, therefore, directly linked to the MTORC1 substrates EIF4EBP1 and RPS6KB1. Upon parameterisation of time course data from MCF-7 or ZR-75-1 cells, the reduced model version matched data very closely (Fig 7). It also matched the simulation where the stable state was determined analytically (see supplement S5). The *χ*^2^ values were calculated for the individual readouts to determine the fit of the simulations to the experimental data for both models (see supplement S1). Depending on the readout the *χ*^2^ values were lower for either with the full model (Fig 6) or with the reduced model (Fig 7). To make a comparison for the overall fit of the two models the Akaike information criterion (AIC) value was calculated (see supplement S1). Here the reduced model showed a better overall fit with an AIC value of 0.0136 compared to the full model that had an AIC value of 0.0168.

**Figure 7.**
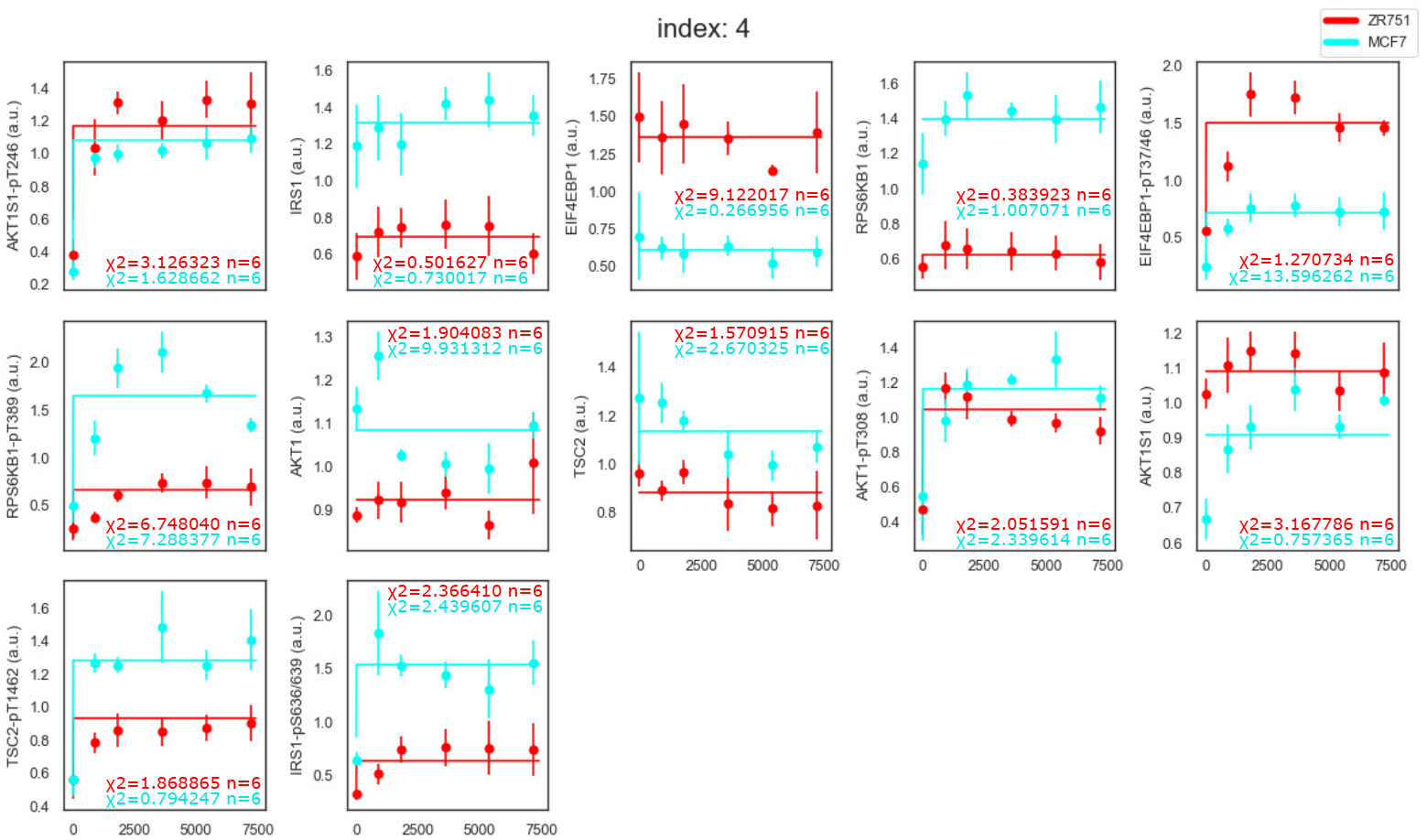
MCF-7 & ZR-75-1 based parameterisation of reduced model. Reduced model parameterised against MCF-7 and ZR-75-1 data (see supplement S2). Parameter set 6 out of sets 0 to 9 which closely matches data (chosen based on custom RSS score). Note that each phospho-protein species (AKT1S1-pT246, AKT1-pS473, AKT1-pT308, TSC2-pT1462, RPS6KB1-pT389, EIF4EBP1-pT37/46 and IRS1-pS636/639) has had a scale factor applied (identical in both cell lines) to the simulation outputs to match them to the experimental data (which has no absolute quantification). Total protein levels (AKT1, AKT1S1, TSC2, RPS6KB1, EIF4EBP1 and IRS1) were not scaled. *χ*^2^ values as described in supplementary S1.4. Dots, experimental data; lines, simulation; bars, standard deviation.

Finally, we sought to leverage the parameterisation acquired by fitting the full model (Fig 3) to MCF-7 data alone to see if it could describe the ZR-75-1 data set. However, the ZR-75-1 data set does not contain data for the initial concentrations of MTOR, RICTOR and RPTOR so an additional parameter estimation was done to estimate these levels assuming the reused parameter set. The result is in close agreement with the ZR-75-1 data set (Fig 8).

**Figure 8.**
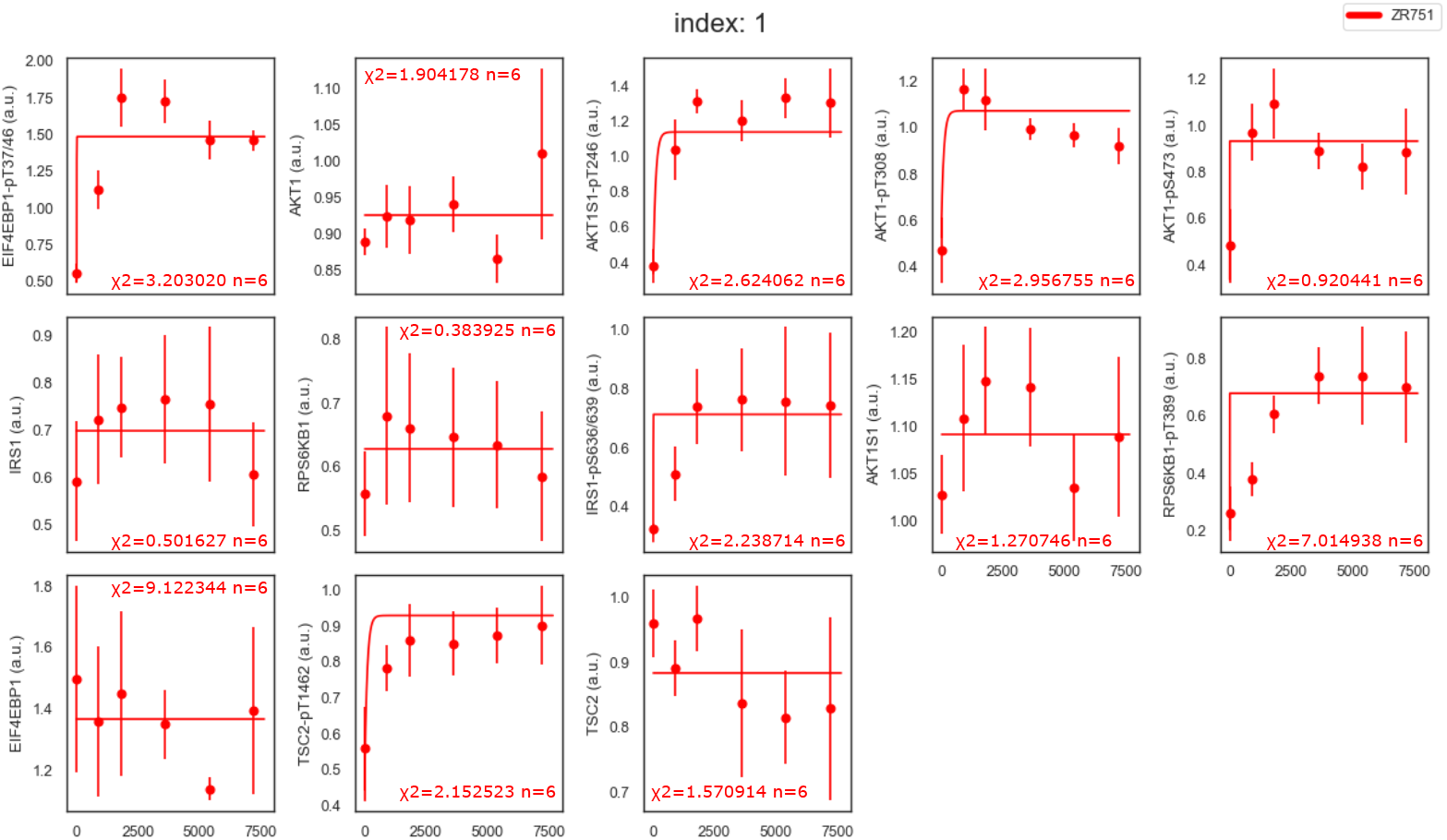
ZR-75-1 behaviour inferred from MCF-7 derived MTOR model. MTOR model using the parameter set from MCF-7 parameterisation (see figure 5) and total protein levels from ZR-75-1 data (see supplement S4). RICTOR, RPTOR and MTOR levels are set using a parameter estimation against ZR-75-1 data. Parameter set 1 out of sets 0 to 9 which closely matches data (chosen based on custom RSS score). Note that each phospho-protein species (AKT1S1-pT246, AKT1-pS473, AKT1-pT308, TSC2-pT1462, RPS6KB1-pT389, EIF4EBP1-pT37/46 and IRS1-pS636/639) has had a scale factor applied to the simulation outputs (based on comparison of the MCF-7 data and its associated simulation) to match them to the experimental data (which has no absolute quantification). Total protein levels (AKT1, AKT1S1, TSC2, RPS6KB1, EIF4EBP1 and IRS1) were not scaled. It should be noted AKT1-pT308/S473 levels contribute to both the AKT1-pS473 and AKT1-pT308 readings. *χ*^2^ values as described in supplementary S1.4. Dots, experimental data; lines, simulation; bars, standard deviation.

The full simulations associated with the full 10 parameter sets can be seen in the supplementary materials.

In summary the technique is able to produce simulations consistent with data while enforcing an initial stable state in multiple models of varied complexity, using semi quantitative data from multiple cell lines. This approach is effective on models that fail to achieve initial stable state with conventional parameterisation.

## Discussion

It is common in experiments on signalling pathways for the appropriate modelling assumption to be that the system starts in a stable state. This can be challenging because models don’t have to become particularly complex with extensive cross talk for analytical calculation of the stable state to become infeasible. Ideally a tool for parameter estimation of systems biology models should have a facility to define a parameter estimation in such a way that initial conditions and unestimated parameters could be set using a pre-simulation of variable length.

It should be noted that other software for semi-quantitative as well as qualitative data are available that implement some of these features. Both, AMICI in combination with pyPESTO [37, 38] and PETab.jl [39, 40] have been released recently and can perform parameter estimation with Pre-Equilibration conditions. However these tools lack a built in system for leveraging the parallelism of a SLURM computer cluster without the user doing additional work. Data 2 dynamics also supports Pre-Equilibration, but is dependent on the proprietary Matlab software [41]. These tools use adaptive strategies for selecting the stopping time to avoid the simulation from getting caught in a limit cycle. COPASI also has a Pre-Equilibration capability but the feature is undocumented and not exposed through the pycotools library. In contrast, our tool for COPASI uses a fixed amount of simulated time during which an equilibrium needs to be reached, or the simulation will be rejected. As our method simulates the ODE system until it stabilizes or until the algorithm decides to reject the parameter set because it takes too long to stabilize, the simulation also will not get caught in limit cycles unless the ODE system has oscillatory behavior.

COPASI has recently acquired its own API ‘basico’. However our tool makes use of the older pycotools library as an API for COPASI. One disadvantage of this is pycotools does not support newer standards like PETab. However a particular advantage of pycotools is that it supports parallel execution on an open source SLURM computer cluster [42, 43]. A number of very recently developed tools such as parPE [44], ParaCopasi [45] and PyBioNetFit [46] also support parameter estimation on a SLURM system. However these tools do not support handling semi quantitive data by normalisation across multiple experiments. Additional coding similar to our approach would be necessary to make these tools compatible with this form of normalisation. Cross experiment normalisation of semi quantitive data is desirable when an estimations parameter space is already large and unidentifiabilities are likely to be an issue if estimated scale factors are introduced.

Our tool fills that niche where there is a need to parameter estimate a model with a large parameter space against semi quantitive data for multiple cell lines / experimental conditions where the relative levels between experiments are meaningful. Particularly where this necessitates parallel computation and estimated scaling factors are unfeasible.

It should be noted that although our results used an older build of COPASI (the default installed on the HPC at the time) it has been tested with COPASI 4.44 using dummy data on a workstation.

Our approach involves user guidance to rearrange and substitute equations to find initial conditions that quickly pre-equilibrate, but even if this step were neglected our method brings pre-equilibration for parameter estimations to pycotools and exposes the utility of the pycotools library to those who do not wish to write custom code.

We therefore suggest that the approach we’ve implemented has wide application for estimating initial conditions. Apart from steady state, our technique could be generalised to support a number of additional ‘start’ conditions that have no steady state including for instance oscillatory systems. This would involve constraining a simulation to start at a particular phase in a steady oscillation and rejecting those parameter sets that do not behave periodically.

Ultimately biochemical processes in cells are stochastic and inherently noisy. True steady states probably only exist approximately in nature. Deterministic models such as the ones we deal with have inherent limitations in addressing such issues. One particular advantage of our process is that parameter sets that stabilise very slowly will be rejected as poor fits, because we pre-equilibrate by running the simulation for a limited period of time which returns high values if the pre-equilibration fails.

The use of semi quantitative data for parameter estimation is widespread. We could have approached the issue of the semi quantitative nature of our data by incorporating estimated scale factors but this would likely have introduced unidentifiabilities and further expanded a large set of estimated parameters. Nor could we easily fix an arbitrary scale since we considered multiple phosphorylation sites of the same protein within our data. This informed our choice of a normalization approach.

In the absence of scale factors most parameterisation software operates under the assumption that model quantities can be compared absolutely with experimental data. To be specific that the simulation output relates to absolute quantities but is compared to semi-quantitative data. We solve this issue by normalizing out absolute quantity from simulated data which allows direct comparison with normalized experimental data. This is only possible if we can modify the outputs of the simulations within the parameter estimation process. It is helpful when parameter estimation software facilitates a post simulation calculation allowing the simulated time courses from a set of multiple simulations to be modified before comparison with experimental data.

On the experimental data side such modifications will of course affect the distribution of error in data. Our approach so far ignores variation between replicates and future work should account for this, considering complex noise distribution [47].

An alternative approach we could have taken to the lack of absolute quantification would be to arrange the parameterisation so that the fold change in species over time is compared to the fold change in data. However, the initial time point for a given species in data may often be very close to zero, resulting in a considerable level of noise relative to the data point value, which introduces uncertainty across the whole simulation time course. This issue is avoided by our approach which normalizes the simulation data across the whole time course.

Another approach would be to treat scale factors as estimated parameters. However, in our case this approach produced challenging parameter estimations that exceeded the available computational resources, likely because of the high number of parameters it added.

Furthermore, in agreement with others [10–12], we have demonstrated that a single model describing multiple cells lines, merely by varying total protein abundances, can be constructed. The technique we’ve developed will be of broad use to facilitate parameter estimation for systems biology models of signalling pathways. We recommend the implementation of variable length pre-simulation periods and post-simulation output calculations in standard parameterisation software.

## Supporting information

S2 data MCF-7 & ZR-75-1

S3 data MCF-7

## Supporting information

**S1 Supporting Information** Technical details regarding data and code as well as equations specifying the models.

**S2 data MCF-7 & ZR-75-1** Quantified experimental data of MCF-7 and ZR-75-1 cells shown in Fig 2 used for model parameterization in Fig 6&7.

**S3 data MCF-7** Subset of quantified experimental data shown in S2: data of MCF-7 cells only. Used for model parameterization in Fig 5.

**S4 data ZR-75-1** Subset of quantified experimental data shown in S2: data of ZR-75-1 cells only. Used for model parameterization in Fig 8.

**S5 Figure** A figure showing a simulation of a model (see Fig 4) parameterized using analytical stabilization of the initial conditions.

## Acknowledgments

The authors would like to thank Kirk Godber for the loan of computing resource used for parts of this work. K.T. acknowledges support from the DFG (German Research Foundation, project No TH 1358/3–2), from the MESI-STRAT project (grant agreement No 754688) which has received funding from the European Union’s Horizon 2020 research and innovation programme, from the European Partnership for the Assessment of Risks from Chemicals PARC (Grant Agreement No 101057014), and from the European Research Council (ERC AdG BEYOND STRESS, grant agreement No 101054429) which have received funding from the European Union’s Horizon Europe research and innovation programme. Views & opinions are those of the authors.

