## Supplementary material for "Collective parameter estimation of related models with an initial stability constraint": S1 Supporting Information

March 25, 2025

#### 1 Github Repository

The code used to generate, parameterise and visualise our model can be found at `https://github.com/npc101-ncl/MESI_PLOS_COMP_BIO`. 'It should be noted that for the tool to run on both the MCF-7 (zr75 data MCF7only.xlsx) and ZR-75-1 (zr75 data ZR75only.xlsx) cell line data a single file (zr75 data.xlsx) is required that includes both datasets. However all data sets were derived from the same set of experiments as indicated by their shared ZR75 prefix.

#### 2 Equations

For reference in Tables S1, S2 and S3 we provide the ODEs behind the model depicted in figure 3.

For simplicity we assumed the volume of the cell is 1. We also provide in Table S4 the ODEs behind the model depicted in figure 4.

#### 3 Prior model attempts using objective function with initial steady state 'cost' term.

An early attempt to ensure models began in a stable state involved rigging COPASI to parameterize both the model with perturbation against dynamic data (fig S1) and without perturbation against a 'steady state' time course (fig S2) constructed by using the first time point as a reference. As can be seen in Fig S2 this approach failed to achieve a steady state consistent with the first time point. This parameterisation uses the data in t47d_data.xlsx.

The files in the GitHub repository used to create these figures were antString5.txt, scaleFinderXParam.py, scaleFinderXPrimer.py, scaleFinderXVisualisor.py, scaleFinderXResults.py, tor5-param.sh and tor5-primer.sh, t47d_data.xlsx. The associated time course data exceeds GitHubs maximum file size. The files have been compressed and can be found in the data/tor5 directory.

Table S1: Part 1 of ODEs for model depicted in figure 3

| variable | expression |
|---|---|
| $\frac{d\,\text{IRS1a}}{dt}$ | Cell * (IRS1 * kIRS1Act * (Insulin + InsulinB) - IRS1a * S6KpT389 * kIRS1Phos) |
| $\frac{d\,\text{IRS1}}{dt}$ | Cell * (-IRS1 * kIRS1Act * (Insulin + InsulinB) + IRS1pS636_639 * kIRS1Inact) |
| $\frac{d\,\text{IRS1pS636\_639}}{dt}$ | Cell * (IRS1a * S6KpT389 * kIRS1Phos - IRS1pS636_639 * kIRS1Inact) |
| $\frac{d\,\text{pPI3K}}{dt}$ | Cell * (IRS1a * PI3K * kPI3KPhos - kPI3KDephos * pPI3K) |
| $\frac{d\,\text{PI3K}}{dt}$ | Cell * (-IRS1a * PI3K * kPI3KPhos + kPI3KDephos * pPI3K) |
| $\frac{d\,\text{AktpT308}}{dt}$ | Cell * (Akt * kAktPhos_kcat * pPI3K - AktpT308 * kAktByTor2Phos * pmTORC2 - AktpT308 * kAktDephos + AktpT308S473 * kAktByTor2Dephos) |
| $\frac{d\,\text{Akt}}{dt}$ | Cell * (-Akt * kAktByTor2Phos * pmTORC2 - Akt * kAktPhos_kcat * pPI3K + AktpS473 * kAktByTor2Dephos + AktpT308 * kAktDephos) |
| $\frac{d\,\text{AktpT308S473}}{dt}$ | Cell * (AktpS473 * kAktPhos_kcat * pPI3K + AktpT308 * kAktByTor2Phos * pmTORC2 - AktpT308S473 * kAktByTor2Dephos - AktpT308S473 * kAktDephos) |

Table S2: Part 2 of ODEs for model depicted in figure 3

| variable | expression |
|---|---|
| $\frac{d\,\mathrm{AktpS473}}{dt}$ | Cell * (Akt * kAktByTor2Phos * pmTORC2 - AktpS473 * kAktByTor2Dephos - AktpS473 * kAktPhos_kcat * pPI3K + AktpT308S473 * kAktDephos) |
| $\frac{d\,\mathrm{TSC2pT1462}}{dt}$ | Cell * (AktpT308 * TSC2 * kTSC2Phos + AktpT308S473 * TSC2 * kTSC2Phos * kTSC2PhosBoost - TSC2pT1462 * kTSC2Dephos) |
| $\frac{d\,\mathrm{TSC2}}{dt}$ | Cell * (-AktpT308 * TSC2 * kTSC2Phos - AktpT308S473 * TSC2 * kTSC2Phos * kTSC2PhosBoost + TSC2pT1462 * kTSC2Dephos) |
| $\frac{d\,\mathrm{mTORC1lys}}{dt}$ | Cell * (TSC2 * kmTORC1Dephos * pmTORC1 - kmTORC1LysToCyt * mTORC1lys - kmTORC1Phos * mTORC1lys - kmTORC1_Dis * mTORC1lys + kmTORC1cytToLys * mTORC1cyt * (AA + AAB)) |
| $\frac{d\,\mathrm{mTORC1cyt}}{dt}$ | Cell * (RPTOR * kmTOR_RPT_Comb * mTOR + kmTORC1LysToCyt * mTORC1lys - kmTORC1_Dis * mTORC1cyt - kmTORC1cytToLys * mTORC1cyt * (AA + AAB)) |
| $\frac{d\,\mathrm{pmTORC1}}{dt}$ | Cell * (-TSC2 * kmTORC1Dephos * pmTORC1 + kmTORC1Phos * mTORC1lys - kmTORC1_Dis * pmTORC1) |
| $\frac{d\,\mathrm{PRAS40pT246}}{dt}$ | Cell * (PRAS40 * kPras40Phos * (AktpT308 + AktpT308S473) - PRAS40pT246 * kPras40Dephos) |
| $\frac{d\,\mathrm{PRAS40}}{dt}$ | Cell * (-PRAS40 * kPras40Phos * (AktpT308 + AktpT308S473) + PRAS40pT246 * kPras40Dephos) |

Table S3: Part 3 of ODEs for model depicted in figure 3

| variable | expression |
|---|---|
| $\frac{d\,\text{FourEBP1pT37\_46}}{dt}$ | Cell * (FourEBP1 * kFourEBP1Phos * pmTORC1 - FourEBP1pT37_46 * kFourEBP1Dephos) |
| $\frac{d\,\text{FourEBP1}}{dt}$ | Cell * (-FourEBP1 * kFourEBP1Phos * pmTORC1 + FourEBP1pT37_46 * kFourEBP1Dephos) |
| $\frac{d\,\text{S6KpT389}}{dt}$ | Cell * (S6K * kS6KPhos * pmTORC1 - S6KpT389 * kS6KDephos) |
| $\frac{d\,\text{S6K}}{dt}$ | Cell * (-S6K * kS6KPhos * pmTORC1 + S6KpT389 * kS6KDephos) |
| $\frac{d\,\text{pmTORC2}}{dt}$ | Cell * (-S6KpT389 * kmTORC2DephosByS6K * pmTORC2 + kRAS * mTORC2 * (Insulin + InsulinB) - kmTORC2Dephos * pmTORC2 - kmTORC2_Dis * pmTORC2) |
| $\frac{d\,\text{mTORC2}}{dt}$ | Cell * (RICTOR * kmTOR_RIC_Comb * mTOR + S6KpT389 * kmTORC2DephosByS6K * pmTORC2 - kRAS * mTORC2 * (Insulin + InsulinB) + kmTORC2Dephos * pmTORC2 - kmTORC2_Dis * mTORC2) |
| $\frac{d\,\text{RICTOR}}{dt}$ | Cell * (-RICTOR * kmTOR_RIC_Comb * mTOR + kmTORC2_Dis * mTORC2 + kmTORC2_Dis * pmTORC2) |
| $\frac{d\,\text{mTOR}}{dt}$ | Cell * (-RICTOR * kmTOR_RIC_Comb * mTOR - RPTOR * kmTOR_RPT_Comb * mTOR + kmTORC1_Dis * mTORC1cyt + kmTORC1_Dis * mTORC1lys + kmTORC1_Dis * pmTORC1 + kmTORC2_Dis * mTORC2 + kmTORC2_Dis * pmTORC2) |
| $\frac{d\,\text{RPTOR}}{dt}$ | Cell * (-RPTOR * kmTOR_RPT_Comb * mTOR + kmTORC1_Dis * mTORC1cyt + kmTORC1_Dis * mTORC1lys + kmTORC1_Dis * pmTORC1) |

Table S4: ODEs for model depicted in figure 4

| variable | expression |
|---|---|
| $\frac{d\,\text{IRS1pS636\_639}}{dt}$ | Cell * (IRS1 * S6KpT389 * kIRS1Phos - IRS1pS636_639 * kIRS1Inact) |
| $\frac{d\,\text{IRS1}}{dt}$ | Cell * (-IRS1 * S6KpT389 * kIRS1Phos + IRS1pS636_639 * kIRS1Inact) |
| $\frac{d\,\text{AktpT308}}{dt}$ | Cell * (Akt * Insulin * kAktPhos_kcat + Akt * kAktPhos_kcatB - AktpT308 * IRS1pS636_639 * kAktDephos) |
| $\frac{d\,\text{Akt}}{dt}$ | Cell * (-Akt * Insulin * kAktPhos_kcat - Akt * kAktPhos_kcatB + AktpT308 * IRS1pS636_639 * kAktDephos) |
| $\frac{d\,\text{TSC2pT1462}}{dt}$ | Cell * (AktpT308 * TSC2 * kTSC2Phos - TSC2pT1462 * kTSC2Dephos) |
| $\frac{d\,\text{TSC2}}{dt}$ | Cell * (-AktpT308 * TSC2 * kTSC2Phos + TSC2pT1462 * kTSC2Dephos) |
| $\frac{d\,\text{PRAS40pT246}}{dt}$ | Cell * (AktpT308 * PRAS40 * kPras40Phos - PRAS40pT246 * kPras40Dephos) |
| $\frac{d\,\text{PRAS40}}{dt}$ | Cell * (-AktpT308 * PRAS40 * kPras40Phos + PRAS40pT246 * kPras40Dephos) |
| $\frac{d\,\text{FourEBP1pT37\_46}}{dt}$ | Cell * (AA * FourEBP1 * kFourEBP1Phos + (FourEBP1 * kFourEBP1PhosB - FourEBP1pT37_46 * kFourEBP1Dephos) * (PRAS40 * kTORPRAS + TSC2 * kTORTSC + 1))/(PRAS40 * kTORPRAS + TSC2 * kTORTSC + 1) |
| $\frac{d\,\text{FourEBP1}}{dt}$ | Cell * (-AA * FourEBP1 * kFourEBP1Phos + (-FourEBP1 * kFourEBP1PhosB + FourEBP1pT37_46 * kFourEBP1Dephos) * (PRAS40 * kTORPRAS + TSC2 * kTORTSC + 1))/(PRAS40 * kTORPRAS + TSC2 * kTORTSC + 1) |
| $\frac{d\,\text{S6KpT389}}{dt}$ | Cell * (AA * S6K * kS6KPhos + (S6K * kS6KPhosB - S6KpT389 * kS6KDephos) * (PRAS40 * kTORPRAS + TSC2 * kTORTSC + 1))/(PRAS40 * kTORPRAS + TSC2 * kTORTSC + 1) |
| $\frac{d\,\text{S6K}}{dt}$ | Cell * (-AA * S6K * kS6KPhos + (-S6K * kS6KPhosB + S6KpT389 * kS6KDephos) * (PRAS40 * kTORPRAS + TSC2 * kTORTSC + 1))/(PRAS40 * kTORPRAS + TSC2 * kTORTSC + 1) |

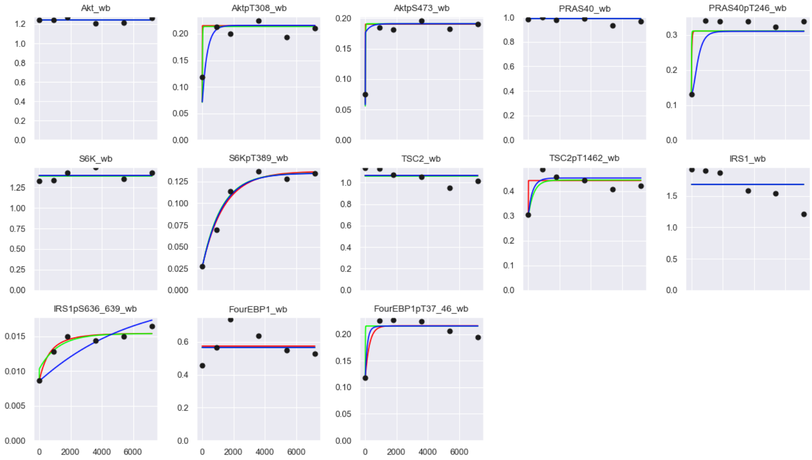


Figure S1: The dynamic (perturbed) time course of the 3 best parameter sets produced by our abandoned approach. MTOR model parameterised against MCF-7 data (see t47d_data.xlsx in the github). It should be noted AKT1-pT308/S473 levels contribute to both the AKT1-pS473 and AKT-pT308 readings. Dots, experimental data; lines, simulation.

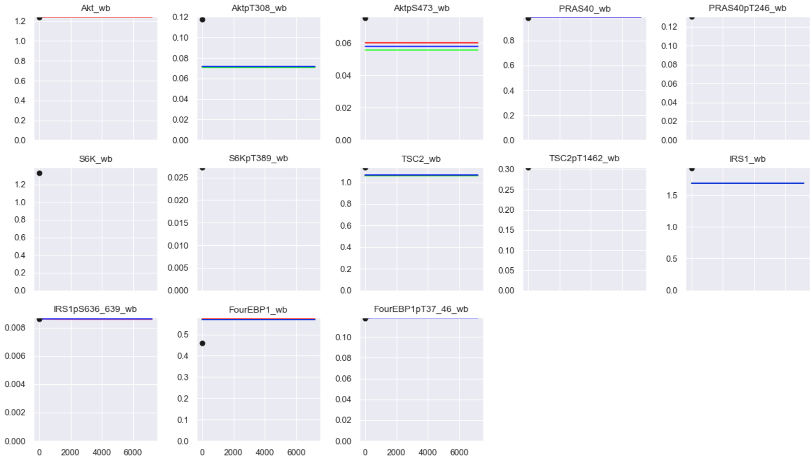


Figure S2: The Stable state (unperturbed) time courses of the 3 best parameter sets produced by our abandoned approach. MTOR model parameterised against MCF-7 data (see t47d_data.xlsx in the github). It should be noted AKT1-pT308/S473 levels contribute to both the AKT1-pS473 and AKT-pT308 readings. Dots, experimental data; lines, simulation.

### 4 Implementation of the chi squared goodness of fit test

Because the outputs of our model are without a relative scale to experimental data it was necessary to adjust the chi squared test. As in fig 5-8 the data given by our model was scaled so the species averages aligned with the mean of the experimental data. Hence the following.

$$\sum_{i,k} y_{ijk} = \sum_{i,k} y'_{ijk},\ \forall\, j$$

Where $i$ is the index over both time points, $k$ cell lines and $j$ the index over observed species. $y_{ijk}$ is an mean observation of the experimental data, $y'_{ijk}$ is the associated scaled model output data. The scaling can be considered analogous to an additional stage of parametrization and so the number of rescalings ($S$) must be considered as additional parameters in the degree of the test. Consequently our chi squared test formula is.

$$\sum_{i} \left( \frac{y_{ijk} - y'_{ijk}}{\sigma_{ijk}} \right)^2 = \chi^2_{n-P-S}$$

Where $P$ is the number of parameters in our original parameter estimation (including estimated initial values where appropriate) and $n$ the number of data points. $\sigma_{ijk}$ is the standard deviation for replicates of a particular data point $y_{ijk}$.

Values of $\chi^2$ and $n$ are listed on figures 5-8. Values generated by 'chiSqGen.py'.

### 5 Sympy adjustment of initial conditions to reduce stabilization time

To validate that our modification of the initial conditions with Sympy derived expressions does in fact reduce the time the model takes to stabilize we performed a set of simulations (see GitHub file stabSpeedTest.py). For a randomly generated set of parameters and total protein levels we simulated 2 version of the model. One with the antStringMSympy.txt antimony string which utilizes sympy to partially solve the ODEs for the stable state and sets some initial conditions and one with antStringMNoSympy.txt which utilizes a best guess based on conservation rules alone. Both antimony strings are derived from the model in figure 3. The graph in fig S3 clearly shows that for a random sample of parameters our method reduces the simulated time to stabilization.

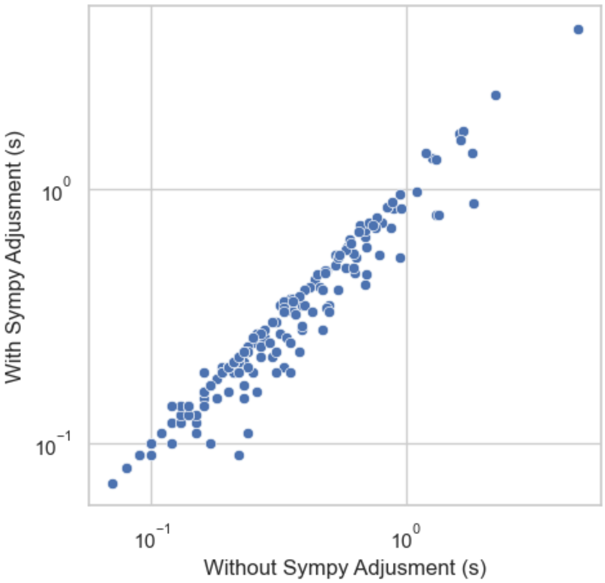


Figure S3: The time in simulated seconds for the model (as described in fig 3) to stabilize with and without adjustment of initial conditions using sympy. The associated data can be found in the GitHub in the sympyTest.csv file. This is a log/log plot.

### 6 Transition Times

Transition times for the model's steady states were computed using the formula

$$T_i = \frac{x_i(t=0)}{\sum_j |R_{i,j}(x_i(t=0))|}$$

Where $T_i$ is the transition time, $x_i$ the species and $R_{ij}$ reactions effecting the species.

Transition times for the model in figure 3 may be found in table S5 and for figure 4 in table S6. Table generated by 'transTimes.py'.

### 7 AIC scores

We computed AIC scores for the models described in figures 5-8 using the following formula.

$$\text{RSS} = \sum_{i=1}^{n} (y_i - y_i')^2$$

$$\text{AIC} = 2k + n \ln\left(\frac{\text{RSS}}{n}\right) + n\ln(2\pi) + n$$

Table S5: transition times (s) for model in figure 3

| Figure<br>Cell Line | 5<br>MCF7 | 6<br>ZR751 | 6<br>MCF7 | 8<br>ZR751 |
|---|---|---|---|---|
| Akt | 1.48e-01 | 5.10e-02 | 3.92e-02 | 1.82e+01 |
| AktpS473 | 3.81e-02 | 5.20e-03 | 5.20e-03 | 4.62e-02 |
| AktpT308 | 2.92e-02 | 5.09e-02 | 3.92e-02 | 1.34e+01 |
| AktpT308S473 | 1.86e-02 | 5.20e-03 | 5.20e-03 | 4.62e-02 |
| FourEBP1 | 2.16e-02 | 1.30e-02 | 9.46e-03 | 2.32e+00 |
| FourEBP1pT37_46 | 1.44e+00 | 1.69e-02 | 1.69e-02 | 1.36e+00 |
| IRS1 | 9.05e+01 | 2.30e-01 | 2.30e-01 | 9.20e+00 |
| IRS1a | 1.07e-02 | 2.64e-02 | 9.69e-03 | 4.79e-01 |
| IRS1pS636_639 | 1.20e+01 | 1.73e-01 | 1.73e-01 | 4.74e+00 |
| PI3K | 1.38e+05 | 3.04e+02 | 4.23e+02 | 6.04e-01 |
| pPI3K | 5.00e+01 | 5.00e-03 | 5.00e-03 | 1.85e-02 |
| PRAS40 | 6.36e+05 | 4.04e+03 | 4.43e+03 | 1.22e+05 |
| PRAS40pT246 | 1.14e+01 | 5.00e+01 | 5.00e+01 | 1.82e+01 |
| RICTOR | 1.89e-03 | 1.94e-03 | 1.84e-03 | 2.45e-04 |
| RPTOR | 3.01e-03 | 1.84e+00 | 2.10e+00 | 1.02e-02 |
| S6K | 4.13e-03 | 1.65e-02 | 1.21e-02 | 2.54e+00 |
| S6KpT389 | 8.06e-02 | 7.50e-03 | 7.50e-03 | 5.71e-02 |
| TSC2 | 6.31e+01 | 4.56e+03 | 4.90e+03 | 3.92e+02 |
| TSC2pT1462 | 1.86e-02 | 5.00e-03 | 5.00e-03 | 9.10e-03 |
| mTOR | 3.78e-03 | 8.10e-03 | 4.95e-03 | 1.34e-02 |
| pmTORC1 | 1.81e-02 | 8.65e-03 | 7.11e-03 | 6.03e-03 |
| mTORC1cyt | 1.30e-03 | 2.04e-01 | 2.00e-01 | 1.07e-02 |
| mTORC1lys | 1.18e-02 | 8.33e-03 | 6.89e-03 | 4.14e-03 |
| mTORC2 | 1.25e-03 | 1.58e-03 | 1.19e-03 | 8.63e-04 |
| pmTORC2 | 8.18e-02 | 2.05e-01 | 8.10e-02 | 1.60e-02 |

Where $y_i$ is the value of a particular replicate and $y'_i$ the associated scaled simulation data point. $n$ is the total number of observations and $k$ the number of estimated parameters (including scalings applied to simulation results). Results may be seen in table S7. Table generated by 'chiSqGen.py'.

### 8 PETab files

To further illustrate the issue faced in the prior work described in section 3 we have created a set of PETab files that loosely replicate our approach to enforcing a pre-perturbation stable state. Unfortunately PETab is not capable of precisely replicating our approach. In particular instead of the approach to scale estimation we took in section 3 we have added scale factors to PETab as estimated parameters. We also added the standard deviation of the data 'noise' to the model as a parameter since there is no way to omit this from a PETab file set.

The related PETab file sets can be found in the PETab folder in our GitHub repository and were created with the PETabGen.py tool.

Table S6: transition times (s) for model in figure 4

| Figure<br>Cell Line | 7<br>ZR751 | 7<br>MCF7 |
|---|---|---|
| Akt | 4.86e+00 | 4.86e+00 |
| AktpT308 | 4.77e+00 | 1.51e+00 |
| FourEBP1 | 1.21e-01 | 1.21e-01 |
| FourEBP1pT37_46 | 2.19e-02 | 2.19e-02 |
| IRS1 | 2.85e-01 | 1.28e-01 |
| IRS1pS636_639 | 1.05e-01 | 1.05e-01 |
| PRAS40 | 1.09e+02 | 1.94e+02 |
| PRAS40pT246 | 5.00e-03 | 5.00e-03 |
| S6K | 1.74e-01 | 1.74e-01 |
| S6KpT389 | 5.00e-03 | 5.00e-03 |
| TSC2 | 1.93e-02 | 3.44e-02 |
| TSC2pT1462 | 5.36e-03 | 5.36e-03 |

Table S7: AIC scores

| Figure | 5 | 6 | 7 | 8 |
|---|---|---|---|---|
| RSS | 2.14e+01 | 3.98e+01 | 3.79e+01 | 1.87e+01 |
| k (param,scale) | 48 (35,13) | 52 (39,13) | 29 (17,12) | 4 (4,0) |
| n | 306 | 612 | 564 | 306 |
| AIC | 1.51e+02 | 1.68e+02 | 1.36e+02 | 2.09e+01 |
