## Supplementary material for "Collective parameter estimation of related models with an initial stability constraint": S5 Figure

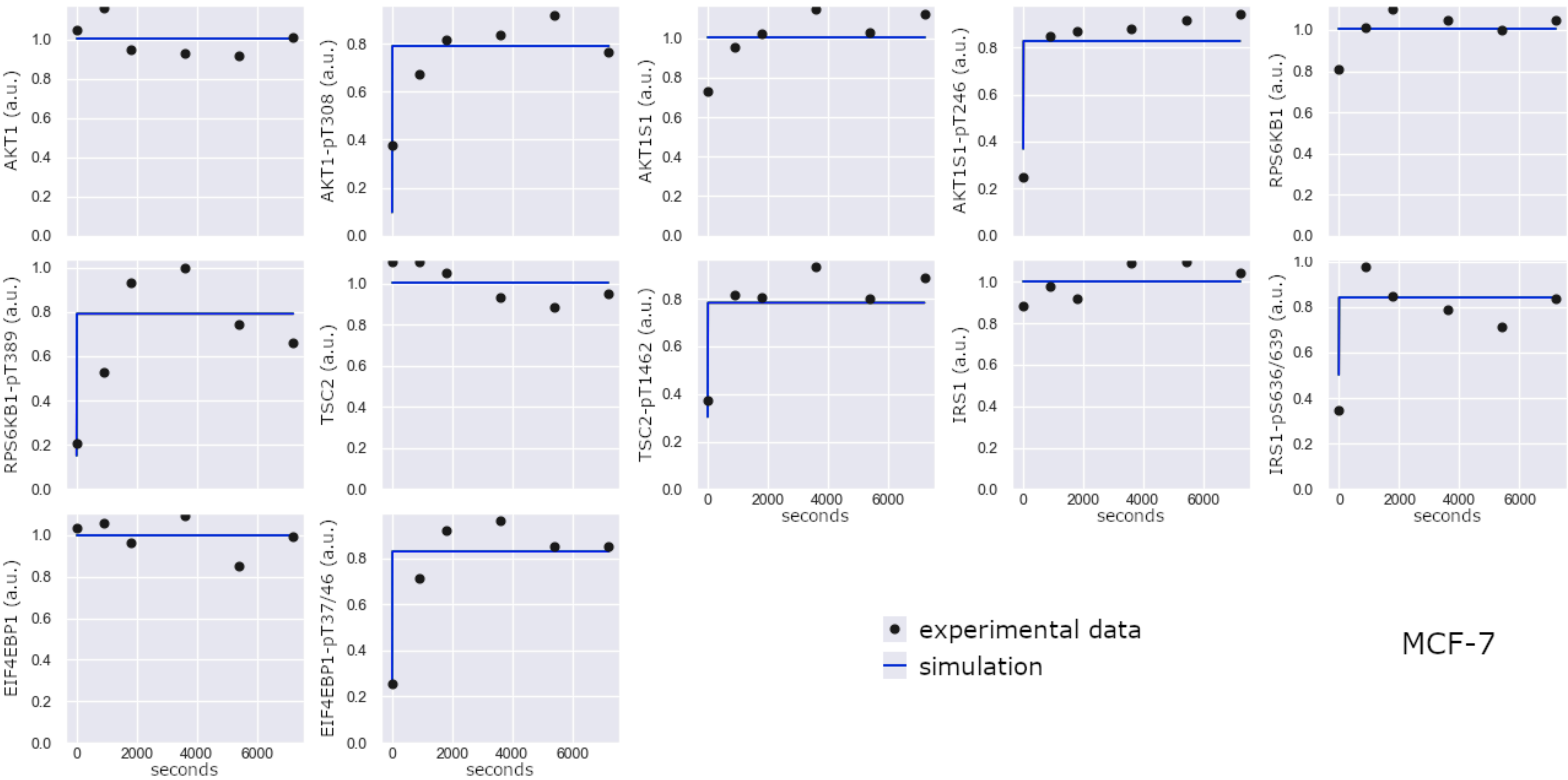


**S5 Figure:** Simulation of model in Fig 4 parameterised using analytical stabilisation of the initial conditions.
